# A teichoic acid-like wall modification associated with immune suppression is socially regulated in *Streptococcus pyogenes*

**DOI:** 10.64898/2026.01.07.698140

**Authors:** Caleb M. Anderson, Reid V. Wilkening, Samy Boulos, Timothy G. Keys, Marc-Olivier Ebert, Léa V. Zinsli, Janes Krusche, Martin J. Loessner, Sam F. Feldstein, Jennifer C. Chang, Andreas Peschel, Alexander R. Horswill, Yang Shen, Michael J. Federle

## Abstract

*Streptococcus pyogenes* (Group A Streptococcus, GAS) is a human-restricted pathogen with a range of clinical manifestations and worldwide prevalence. The GAS Rgg2/Rgg3 quorum sensing (QS) system, a cell-to-cell communication network, modifies the cell surface resulting in increased lysozyme resistance, biofilm formation, and expression of the *qim* operon that is responsible for modulation of innate immune responses in macrophages. The operon encodes 10 genes with predicted homology to enzymes involved in bacterial cell surface-associated carbohydrate and teichoic acid biosynthesis pathways. Comparing extracts of GAS cell wall polysaccharides between wildtype and operon mutants determined that the QS-induced genes modify the *S. pyogenes* cell surface by adding a wall teichoic acid-like moiety of *N*-acetylglucosamine-linked ribitol (GlcNAc-Rbo). A fluorescently labeled phage receptor-binding protein, RBP-13-GFP, that recognizes GlcNAc-decorated ribitol phosphate repeats, bound to the GAS surface only when *qim* expression was induced. Deletion of the *qim* operon eliminated RBP-13-GFP binding, diminished bacterial colonization, and significantly attenuated GAS pathogenesis in a murine skin infection model. These findings indicate that GAS has evolved a strategy to evade innate immune response by presenting a previously unknown carbohydrate moiety upon quorum sensing.

**IMPORTANCE:** *Streptococcus pyogenes* is a major human pathogen, responsible for diverse clinical manifestations of both superficial and invasive infections and can lead to post-infection sequelae like rheumatic heart disease whose prevalence on a global scale rivals the most serious pathogens. Invasive *S. pyogenes* infections are currently on the rise worldwide, notably correlating with increasing pediatric cases of scarlet fever and enhancing the concern for long term complications. There is much that remains unknown about *S. pyogenes* virulence and pathogenicity, and studies focused on understanding basic systems regulating virulence factors could lead to better therapeutics and translational research. We show here one such example, where a bacterial communication system regulating a virulence mechanism relevant to *in vivo* infection confers the ability to alter the host innate immune response. We find that modifications to the cell wall arise when this virulence system is activated that has a direct role in host-pathogen interactions. Further research into this system could provide a mechanism for disruption and serve to treat *S. pyogenes* infection.

## INTRODUCTION

*Streptococcus pyogenes*, also referred to as Group A Streptococcus (GAS), is one of the longest-studied microorganisms, yet our understanding of its numerous virulence factors and role in a diverse range of diseases states remains significantly understudied compared to other human pathogens ^1–6^. GAS is commonly known by its clinical manifestation of streptococcal pharyngitis (strep throat) but is also responsible for diseases such as rheumatic heart disease, scarlet fever, impetigo, necrotizing fasciitis, and streptococcal toxic shock syndrome (STSS). This is reflected in an annual worldwide burden of over 700 million cases of GAS infection, with more than 18 million of those classified as severe disease, together resulting in over 500,000 deaths each year ^3–5^. A rise in invasive GAS infections has recently been observed in the United States and Europe, specifically in the prevalence of scarlet fever and invasive GAS infections in pediatric patients^7^. This is especially concerning given the long-term associated morbidity with the immune sequelae of GAS infections, which far surpasses that of most severe human pathogens^3–5^.

Contributing to the etiology of such a wide range of diseases, the genome of GAS encodes many virulence factors that enable resistance and evasion of host immune responses^8^. Some of these factors are controlled by multifaceted regulatory systems that incorporate cell-to-cell communication networks employing extracellular signaling molecules (pheromones) that facilitate coordination of gene expression in a process known as quorum sensing (QS)^9^. Although several QS systems have been described for GAS, their implication in host immune responses and infection has yet to be fully characterized^10^.

The Rgg2/Rgg3 QS system functions to control transcription of genes at distinct genetic loci: *stcA* (*spy49_0414c*) and the *qim* (*spy49_0450-0460)* operon (Supplement Fig. S1A)^11^. Two Rgg (**r**egulator **g**ene of **g**lucosyltransferase) transcriptional regulators with opposing functions control expression of the regulons, with Rgg2 serving as the transcriptional activator of this system and Rgg3 as a repressor^11–13^. Upon binding of the cognate peptide pheromones (<u>s</u>hort <u>h</u>ydrophobic <u>p</u>eptides, i.e.,SHPs) to the transcriptional regulators, Rgg3 derepresses and allows Rgg2 to activate expression of target genes. Included within the regulon controlled by Rgg2/Rgg3 are the *shp* pheromone genes themselves, generating a positive feedback loop that results in rapid activation of the system once SHP peptides reach nanomolar concentrations^14, 15^.

Previous work examined the role of the Rgg2/Rgg3 QS system on the innate immune response during *in vitro* macrophage infection and found that Rgg2/Rgg3 QS-ON GAS actively suppressed the innate immune response in macrophages in a host-pathogen contact-dependent manner^16, 17^. This infection outcome contrasted with that of QS-OFF GAS, which robustly activated the NF-κB response in a macrophage reporter assay. Positioned downstream of *shp2* and induced in the QS-ON state, *stcA* encodes a positively charged polypeptide that associates with the bacterial cell wall and confers increased lysozyme resistance and biofilm formation^18^. However, the immunomodulatory phenotype did not require *stcA*, but instead was found to be dependent on the *spy49_0450-0460* operon, which is located downstream of *shp3* and also induced by Rgg2 in the QS-ON state^17^.

The *spy49_0450-0460* operon is conserved in all fully sequenced GAS genomes, but very little is understood in how these genes contribute to GAS fitness^19^. Only *spy49_0459*, a HasB paralog annotated as HasB2, has been functionally characterized and found to have UDP-glucose dehydrogenase activity, partially complementing a *hasB* mutant in restoring capsule production^20^. A signature-tagged transposon screen identified mutants of *spy49_0459* and *spy49_0460* to be attenuated for virulence in an invasive zebrafish model, but by undetermined mechanisms^21^. Finally, we recently reported that expression of *spy49_0460* affects the production of several surface-associated virulence factors as well as QS-induced susceptibility to aminoglycosides^22^.

GAS contains multiple surface-associated polysaccharides (Supplemental Fig. S1B) implicated in virulence and pathogenicity with the major known structures in GAS being a hyaluronic acid (HA) capsule, lipoteichoic acid (LTA), and the Group A-specific Carbohydrate also known as GAC^23–35^. Though *hasB2* (*spy49_0459*) can substitute for *hasB*, deleting *hasAB* eliminates HA capsule production altogether, and this deletion did not disable the QS-dependent immunomodulatory phenotype during *in vitro* macrophage infection^17^. Therefore, capsule does not appear to be responsible for the immunosuppressive phenotype and was not evaluated in the current study. LTA in GAS is produced as a polyglycerol-phosphate (polyGroP) teichoic acid, can be decorated with alanine side moieties, and has known immunostimulatory properties with variations in the linkage of LTA to its lipid anchor found to elicit differential host immune responses^31, 32, 36–38^. Many bacterial species also produce peptidoglycan (PG)-bound wall teichoic acids (WTA) and secondary cell wall polysaccharides (SCWP), and while no WTA moieties have been described for GAS^39, 40^, the major SCWP is GAC, a polyrhamnose (polyRha)-based carbohydrate with *N*-acetylglucosamine (GlcNAc) side chains^28, 30, 41, 42^. Approximately 25% of these GlcNAc side chains are further decorated with glycerol phosphate (GroP), which modulates GAS susceptibility to Zn^2+^ toxicity and human antimicrobials (hGIIA and LL-37) ^43^. Teichoic acids and surface-associated carbohydrates are known pathogen-associated molecular patterns (PAMPs) recognized by the host immune system to trigger the innate immune response, and these factors have repeatedly been demonstrated to play a role in bacterial virulence and pathogenicity ^35, 44–51^.

Here, we provide evidence that signaling by the Rgg2/Rgg3 QS system in GAS leads to a modification of the cell wall that correlates with immunosuppression and virulence during *in vivo* colonization. We show that genes of the *spy49_0450-0460* operon (named *qim, for **q**uorum-regulated **i**mmunomodulatory **m**odification*) are required for the wall modification and contributes to GAS virulence during skin colonization. We propose these modifications are responsible for the contact-dependent altered innate immune response observed during *in vitro* macrophage infection.

## RESULTS

### Innate immune suppression requires the *qim* operon and is modulated by genes that affect surface electrostatics

Under culture conditions using a chemically defined, nutrient-rich medium, the Rgg2/Rgg3 system remains in an inactive state due to pheromone turnover by the PepO endopeptidase causing SHP concentrations to remain at sub-stimulatory concentrations; we refer to this state as “QS-OFF”. However, upon the addition of SHP peptide, cultures are rapidly stimulated, leading to robust activation of the QS system within minutes; herein, we refer to the stimulated state as “QS-ON” ^11, 52^. The role of various GAS surface-associated polymers in activation of innate immune responses was monitored using the RAW-Blue macrophage NF-κB reporter bioassay. Rahbari *et al.* ^17^ previously demonstrated that QS-OFF cells induce a high NF-κB response that is enhanced further when co-treated with toll-like receptor (TLR) agonists like lipopolysaccharide (LPS), a well-studied PAMP that is not endogenous to GAS. However, QS-ON bacteria (stimulated with SHP, Fig. 1A) suppress NFκB activation but require the *spy49_0450-0460* (*qim*) operon; likewise, suppression is not an effect of SHP itself as reported previously (Fig. 1B). Repression was restored to the *qim* deletion mutant (Δ*qim*) upon genetic complementation when the operon was placed on a chromosomally integrating plasmid under a constitutive promoter (Fig. 1B). Removal of individual genes from the complementation plasmid, generating operons lacking *spy49_0450* (*qimA*), *spy49_0451* (*qimB*), *spy49_0455* (*qimE*), *spy49_0456* (*qimF*), *spy49_0457* (*qimG*), or *spy49_0458* (*qimH*), were unable to suppress the innate immune response; thus, genetic disruption of any of these genes individually abolished the immunomodulatory ability (Fig. 1C). Prior tests of *spy49_0459* (*qimI*) and *spy49_0460* (*qimJ*) determined that *qimI*, and not *qimJ*, contributed to the suppressive phenotype^17^, whereas *qimC* and *qimD* deletions are planned for future studies.

**Figure 1.**
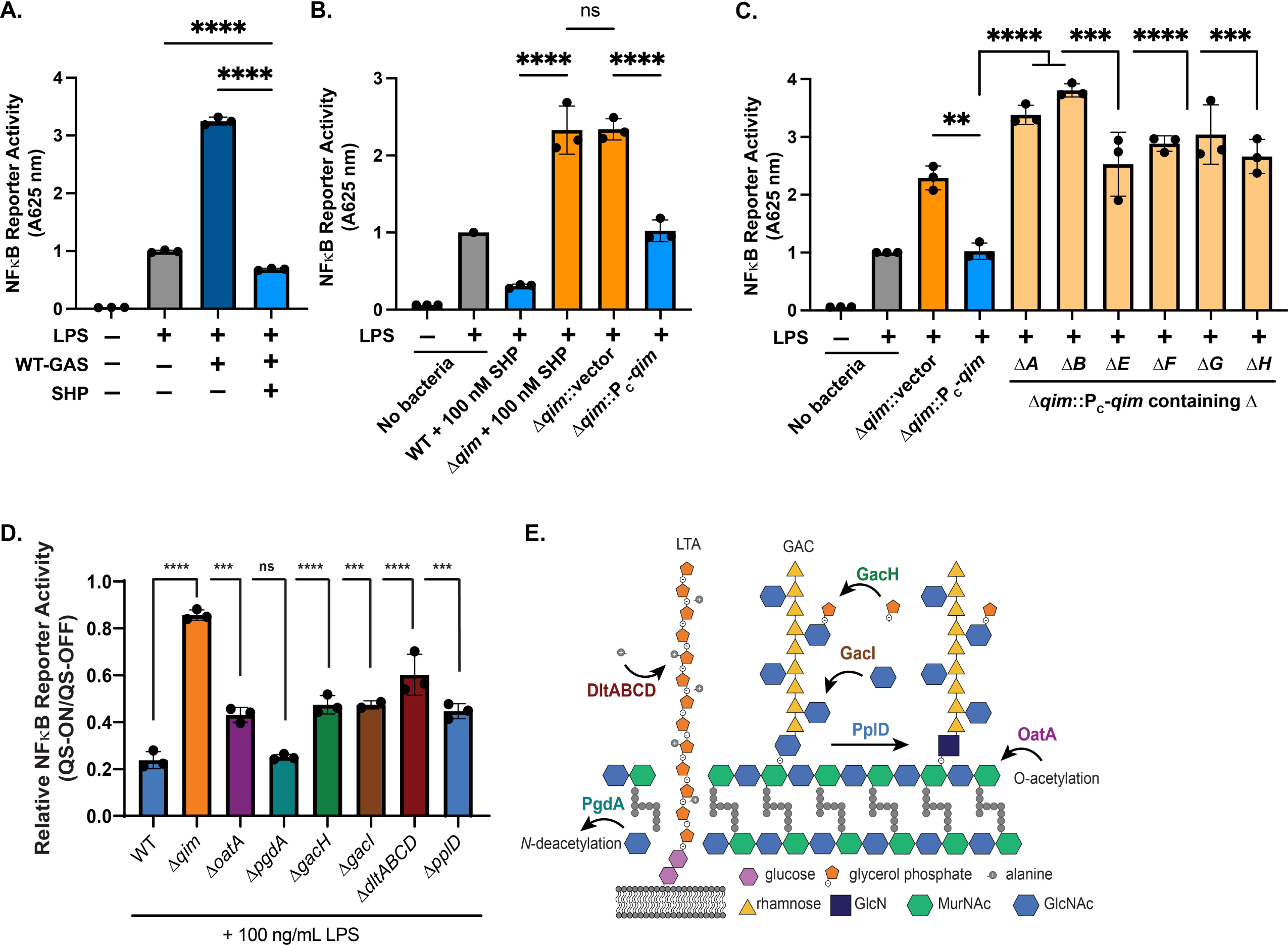
QS-ON GAS alters macrophage innate immune response to infection dependent on the *qim* operon (*Spy49_0450-0460*). **(A)** RAW-Blue^TM^ macrophages contain a secreted embryonic alkaline phosphatase that reports activation of NF-κB and was used for GAS infections *in vitro*. Stimulation of NF-κB was determined for macrophages infected with wildtype (WT) GAS that had been cultured with (QS-ON) or without (QS-OFF) 100 nM SHP. Where indicated, macrophages were stimulated with 100 ng/ml LPS. **(B)** NF-κB reporter assay of macrophages infected with wildtype GAS (+100 nM SHP, QS-ON), Δ*qim* (+100 nM SHP, QS-ON), Δ*qim* with empty vector control (Δ*qim*::p7INT), or Δ*qim* with the *qim* operon expressed under a constitutive promoter on chromosomally integrated p7INT (Δ*qim*::P_C_-*qim*). Results are from three independent experiments conducted in triplicate with values normalized to LPS-only stimulation. **(C)** NF-κB reporter assay of LPS-stimulated macrophages infected with Δ*qim,* Δ*qim* complemented with the complete operon (Δ*qim*::P_C_-*qim*), or Δ*qim* complemented with the operon containing single gene deletions (Δ*A*–Δ*I*). Results shown are from three independent experiments, each conducted in triplicate and normalized to LPS-only treated macrophages. **(D)** NF-κB reporter assay of LPS-stimulated macrophages infected with GAS isogenic mutants lacking cell wall modifying genes: *oatA* (MurNAc *O*-acetylation); *pgdA* (GlcNAc *N*-deacetylation of peptidoglycan), *gacH* (glycerol phosphate modification on GAC); *gacI* (GlcNAc modification on GAC); *dltABCD* (alanylation of lipoteichoic acid, LTA); or *pplD* (GlcNAc *N*-deacetylation of Group A Carbohydrate, GAC). Displayed are the ratios of values resulting from the NF-κB reporter in which macrophages were infected with bacteria treated with 100 nM SHP (QS-ON) vs 0 nM SHP (QS-OFF). **(E)** Schematic indicating cell wall modifications of each gene tested. For panels A-D, means plus standard deviations (SD) are shown with statistical significance indicated as follows: **, *P* < 0.005; ***, *P* < 0.001; ****, *P* < 0.0001 with two-tailed unpaired *t* test (A) and ordinary one-way ANOVA with Tukey’s multiple-comparison test (B, C, and D); ns, no statistical significance.

The degree of QS-dependent suppression was also evaluated and compared between isogenic mutant strains lacking various known cell wall-modifying enzymes. Macrophages were infected with GAS cultures treated with a DMSO-vehicle control (QS-OFF) or treated with 100 nM SHP (QS-ON) for strains lacking the following genes: *oatA* (*spy49_0035*), *pgdA* (*spy49_1092c*), *gacH* (*spy49_0619*), *pplD* (*spy49_0642*), or the *dltABCD* operon (*spy49_1034c-1037c*) (Fig. 1D and 1E). OatA is responsible for *O*-acetylation of *N*-acetylmuramic acid (MurNAc) in the peptidoglycan backbone, while PgdA functions to *N*-deacetylate *N*-acetylglucosamine (GlcNAc) on peptidoglycan ^53–56^. GacH produces a glycerol phosphate (GroP) modification to the GlcNAc side chain of Group A Carbohydrate (GAC) and PplD is a secondary *N*-deacetylase targeting the GlcNAc linkage of GAC to peptidoglycan^43, 57^. The *dlt* operon is responsible for alanylation of lipoteichoic acid ^38^. Each of these enzymes are responsible for modifications that impact electrostatic interactions among structures on the bacterial cell wall. None of the mutants fully abrogated the immunosuppressive phenotype seen for QS-ON GAS infected macrophages but modest reductions could be seen in a gene-specific manner (Fig. 1D). Together these results indicate that the *qim* operon produces an independent mechanism for suppression of the innate immune response; however, modifications to the cell wall electrostatic potential affect the degree to which QS-ON GAS can modulate the innate immune response, which we hypothesize could be due to altered interactions between surface-associated structures. Additionally, all genes evaluated within the *qim* operon are implicated in the biosynthesis of this suppressive factor and required for wildtype level suppression to occur.

### *The qim* operon is critical for virulence in a murine skin colonization model

To evaluate the impact that the Rgg2/Rgg3 regulon provides to bacterial virulence, a murine skin infection model was employed^58^ (62). Wildtype GAS and isogenic mutants of the two primary Rgg2/Rgg3 target loci, Δ*stcA* and Δ*qim*, were each applied to mice individually (Fig. 2A) and over the course of 7 days, animal weights were monitored and their backs swabbed to sample CFU counts present on the skin. By the third day, and continuing through the remainder of the experiment, a significant difference in weights of mice was seen between the uninoculated control group compared to wildtype- or Δ*stcA* mutant-inoculated mice (Fig. 2B). However, no statistical difference of mean weights was seen between the control and Δ*qim-*inoculated groups. By seven days post infection, significant differences were also observed in bacterial burdens as observed by CFU counts after swabbing the treated area (Fig. 2C). Lower amounts of CFUs were recovered from mice treated with both Δ*qim* and Δ*stcA* mutants as compared to wildtype, although a much greater loss of bacteria was seen for the Δ*qim* mutant group. Striking visual differences in infection severity were also observed between the strains, where the Δ*qim* infection resembled the PBS control treatment and Δ*stcA* infection visually mimicked infection with wildtype GAS (Fig. 2D, Supplemental Figs. S2-S5). These results highlight the role that the *qim* operon plays in GAS virulence during skin colonization and validates previous *in vitro* findings that innate immune cell function is compromised when the Rgg2/Rgg3 QS system, and specifically this operon, is active.

**Figure 2.**
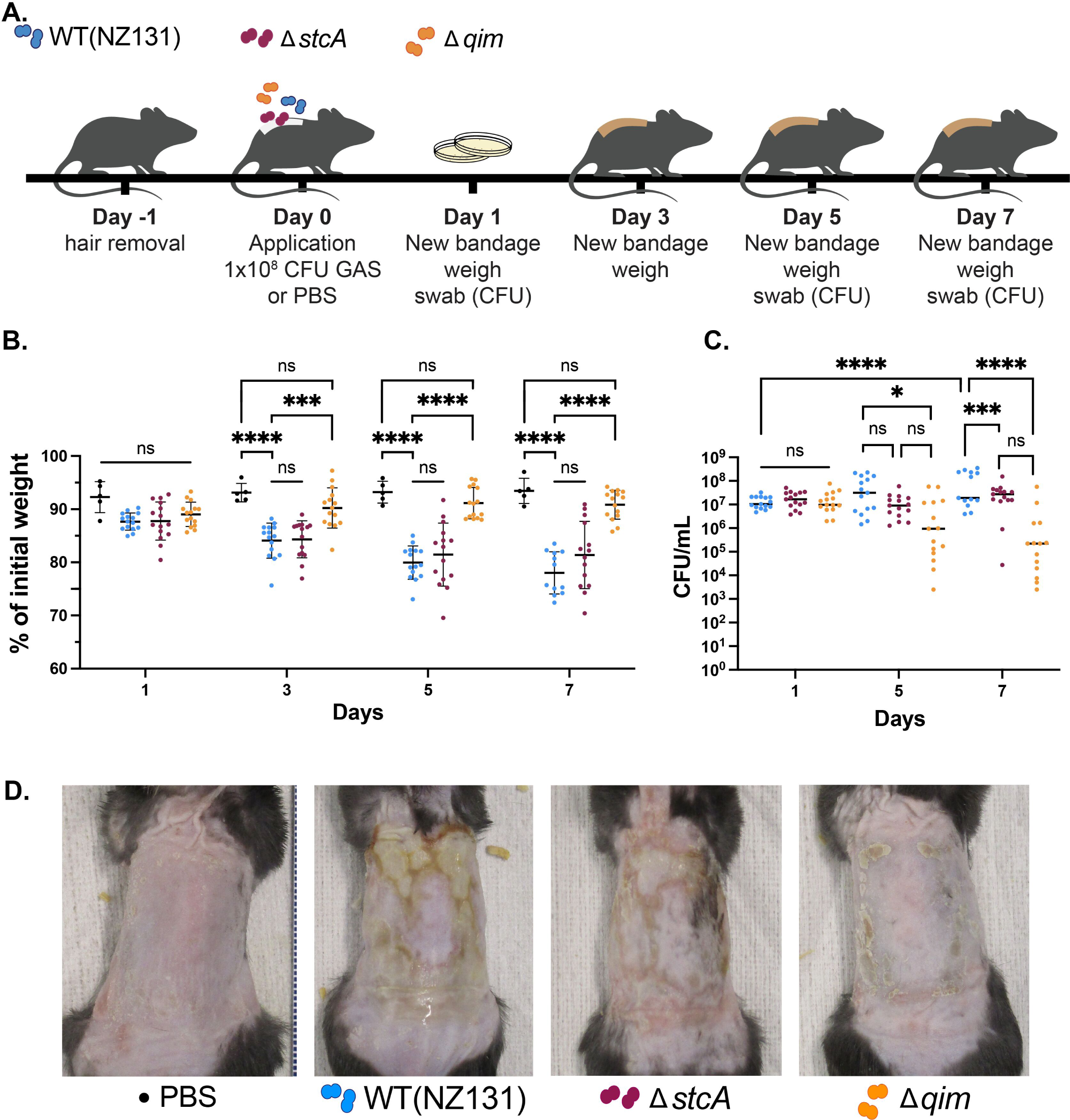
Deletion of the Rgg2/Rgg3 QS-regulated *qim* operon diminishes virulence during murine skin colonization. **(A)** Murine skin colonization model timeline for mice exposed to wildtype and isogenic Δ*stcA* or Δ*qim* mutants. **(B)** Weight loss expressed as percent (%) of initial weight at days 1, 3, 5, and 7 post infection. **(C)** Bacterial burdens as CFU/ml obtained from skin swabs at days 1, 5, and 7 post infection. Results of PBS control group (n=5) and treatment groups (n=15/strain) indicate median (horizontal line) and standard deviations (bars), with statistical significance indicated as follows: *, *P* < 0.05; ***, *P* < 0.001; ****, *P* < 0.0001 by ordinary one-way ANOVA with Tukey’s multiple-comparison test; ns, no statistical significance. (D) Representative images of mouse skin conditions at Day 7. High resolution images of all mice is available in supplementary data.

### *qim* genes share homology to carbohydrate biosynthesis pathways and alter the cell wall-associated carbohydrate content

Initial screening to elucidate the immunomodulatory factor produced by the *qim* operon followed classical bioactivity-guided fractionation using the RAW-Blue macrophage NF-κB reporter bioassay. However, none of the QS-ON GAS extracts generated from solvents covering a range of polarities were able to suppress macrophage stimulation by lipopolysaccharide (LPS) (Fig. 3A). Bioinformatic analysis of the genes contained in the operon revealed homology to enzymes involved in the biosynthesis of cell surface-associated carbohydrates or polymers, indicating the operon’s product is likely not a freely extractable metabolite (Table 1). Structural predictions for each gene were produced using AlphaFold 2.0 and the predicted protein structures were aligned to the top hits generated by the HHpred tool which searches for protein homology (Supplement Fig. S6) ^59–61^. Notably genes *spy49_0451* (*qimB*), *spy49_0454* (*qimD*), *spy49_0457* (*qimG*), *spy49_0459* (*qimI*), and *spy49_0460* (*qimJ*) were all found to have homology to enzymes involved in the biosynthesis of lipopolysaccharide, lipoteichoic acid, wall teichoic acid, and hyaluronic acid capsule in various bacterial species.

**Figure 3.**
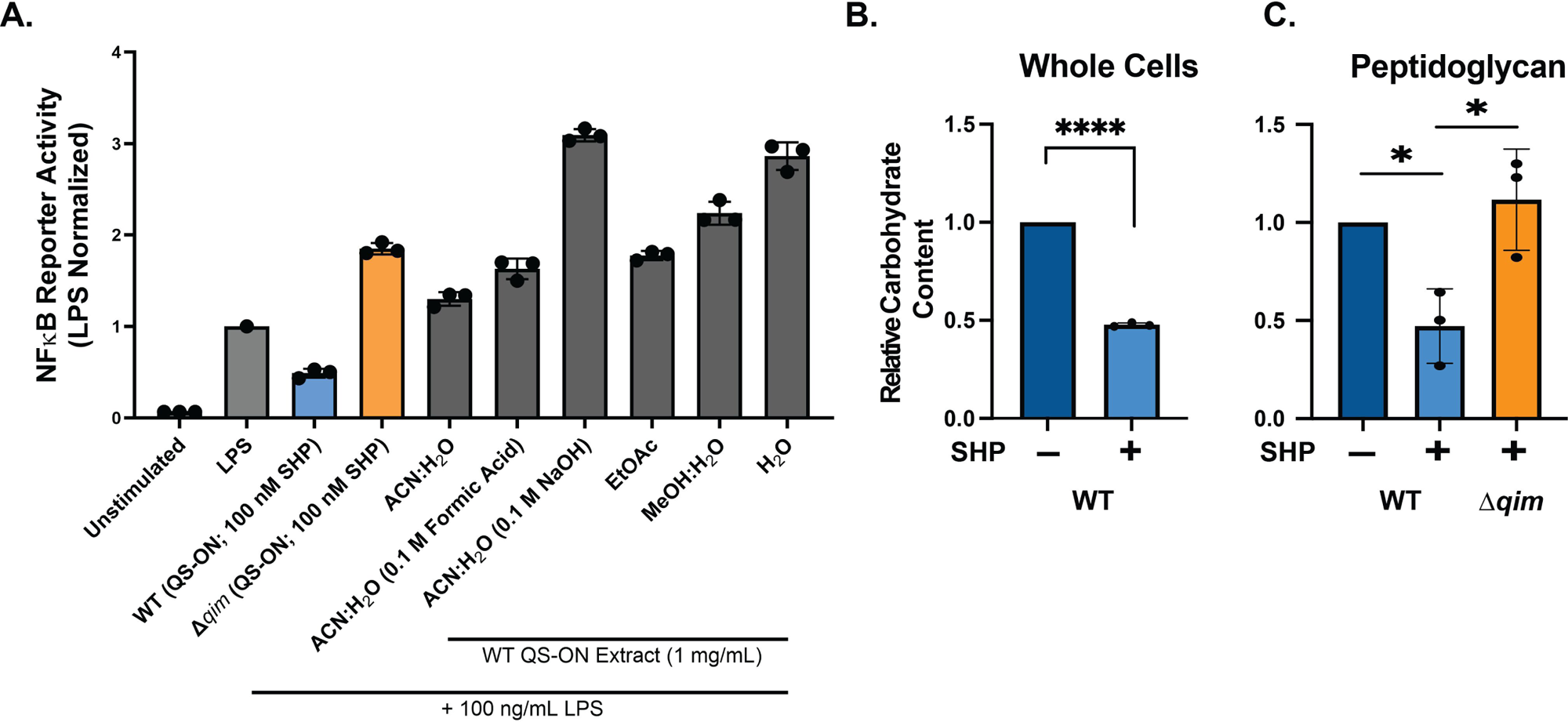
**(A)** Macrophage NF-κB reporter assay of macrophages exposed to extracts obtained from QS-ON GAS. All extracts were provided at 1 mg/mL in macrophage culture medium (DMEM) and results (black bars) normalized to macrophages stimulated with LPS only. Abbreviations: acetonitrile, ACN; ethyl acetate, EtOAc; methanol, MeOH. For comparison, macrophages were infected with wildtype and Δ*qim* that were stimulated with 100 nM SHP (QS-ON). Total carbohydrate content determined by anthrone assay of whole bacterial cells **(B)** or of isolated sacculi **(C)** from wildtype and Δ*qim* cells cultured without (QS-OFF) or with (QS-ON) 100 nM SHP. Results are from three biological replicates; statistical significance is indicated as follows: *, *P* < 0.05; ***, *P* < 0.001; ****, *P* < 0.0001 by ordinary one-way ANOVA with Tukey’s multiple-comparison test.

**Table 1.**
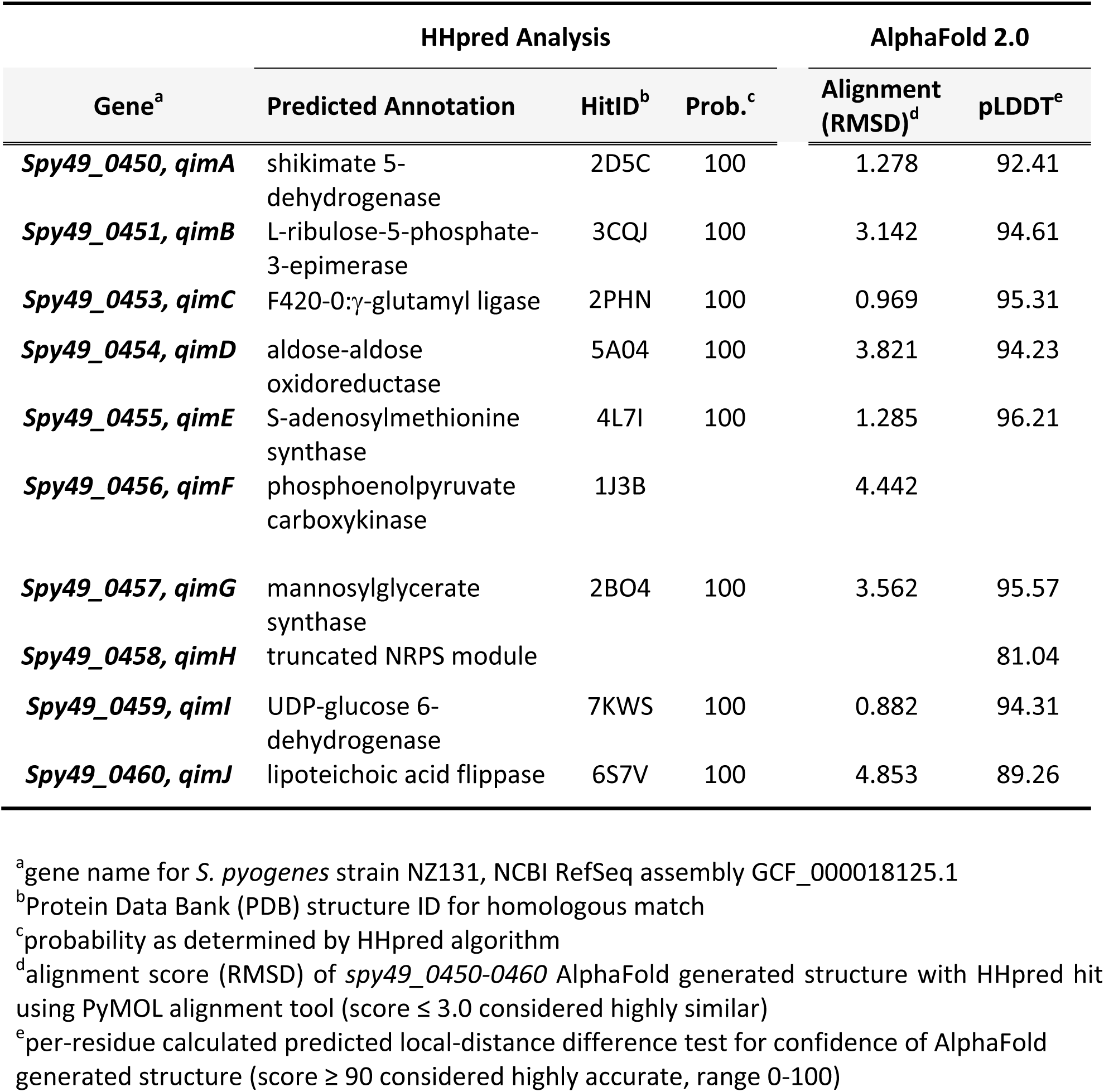
*qim* operon bioinformatic analysis by HHpred and AlphaFold 2.0.

To assess overall amounts of carbohydrate produced by cultures of wildtype QS-OFF and QS-ON we utilized anthrone reagent that quantifies carbohydrates in free and conjugated forms after acid hydrolysis ^62^. Surprisingly, the amount of carbohydrate measured from whole cells was diminished approximately 2-fold when the Rgg2/Rgg3 system was in the QS-ON state for 3 generations over approximately 2 hours (Fig. 3B). The anthrone assay was also conducted on extracted peptidoglycan from wildtype (QS-ON and QS-OFF conditions) and the QS-ON Δ*qim* strain to indicate the overall amount of carbohydrate within or attached to peptidoglycan. This analysis revealed that the QS-dependent reduction of glycans required the presence of *qim*, as the deletion mutant produced similar amounts of carbohydrates as QS-OFF wildtype cells (Fig. 3C). Together with the bioinformatic analyses, these results support the notion that the *qim* operon affects carbohydrate biosynthesis or the modification of polysaccharide(s) attached to or associated with the bacterial cell surface.

### Neither GlcN-linked GAC nor LTA are affected by the *qim* operon

Group A Carbohydrate (GAC) makes up a significant portion of the bacterial cell wall of GAS and is covalently attached at the C6 position of MurNAc of peptidoglycan via a GlcNAc phosphate linkage unit^39, 57^. The *N*-deacetylase PplD converts approximately 80% of these linkage units to glucosamine (GlcN)^57^. This modified linkage unit has increased acid stability which requires extraction by nitrous acid (HONO) for GAC isolation^57^. We compared GlcN-linked GAC from QS-ON GAS to the QS-ON-Δ*qim* mutant by performing fluorescently-labeled glycan compositional analysis (Fig. 4A)^63, 64^. This analysis confirmed the composition of GlcN-linked GAC contains GlcNAc and rhamnose (Rha) carbohydrates; however, no differences were observed between the wildtype and mutant strains. Further characterization by 1D and 2D NMR experiments also revealed no apparent differences between strains and results were nearly identical to previously published structural data for Group A Carbohydrate (Supplemental Table 3, Supplemental Fig. S7) ^43, 57^.

**Figure 4.**
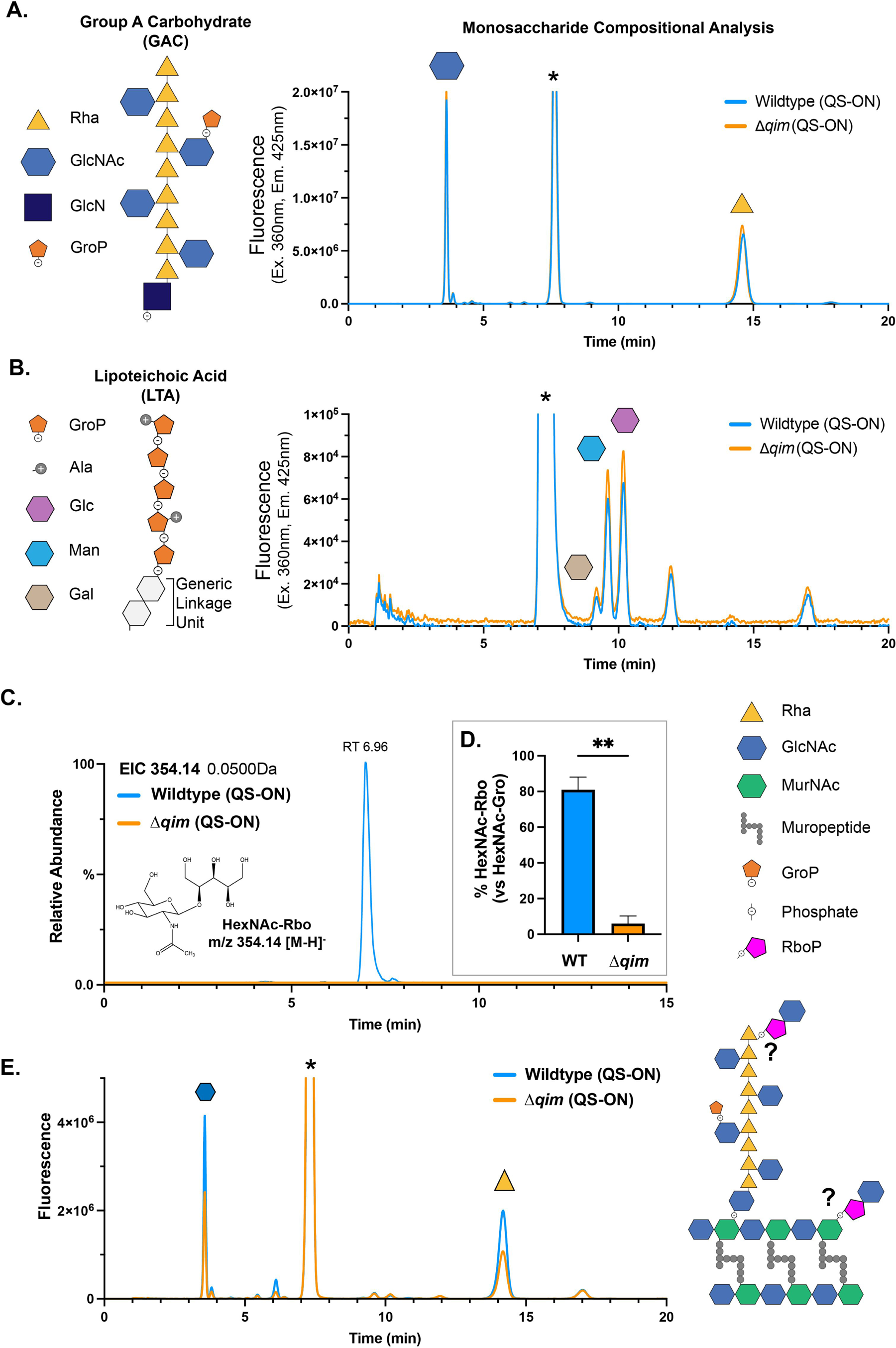
Glycan analysis of GAS polysaccharides identifies RboP-GlcNAc. **(A)** 2-anthranilic acid (2-AA) labeled glycan monosaccharide compositional analysis of GAC linked to peptidoglycan via a GlcN phosphate linker extracted from wildtype and Δ*qim* cultures treated with 100 nM SHP (QS-ON). Chromatogram shows sugars with reducing ends labelled by 2-AA after total acid hydrolysis, separated and detected by HPLC-FLD (excitation 360 nm, emission 425 nm). Abbreviations: Rhamnose, Rha; *N*-acetylglucosamine, GlcNAc; glucosamine, GlcN; glycerol phosphate, GroP. **(B)** 2-AA labeled glycan monosaccharide compositional analysis after total acid hydrolysis of lipoteichoic acid (LTA) extracted from wildtype and Δ*qim* cultures treated with 100 nM SHP (QS-ON). Sugars with reducing ends labelled by 2-AA are shown after separation by HPLC-FLD. **\*** denotes free 2-AA reagent. **(C)** UHPLC-MS/MS extracted ion chromatogram (EIC) of *m/z* 354.14 corresponding to HexNAc ribitol phosphate from mild acid extracted and hydrofluoric acid digested carbohydrate from wildtype QS-ON GAS. The structure of HexNAc-Rbo is shown for reference, with a proposed C2 or C4-linkage via phosphate (not shown). **(D)** Relative percentage of GlcNAc-Rbo to GlcNAc-Gro from isolated sacculi of wildtype and Δ*qim* cells grown under QS-ON conditions (100 nM SHP). Results shown are from two biological replicates (n=2) with means plus standard deviations (SD). **(E)** 2-AA labeled glycan monosaccharide compositional analysis of mild acid extracted carbohydrate from wildtype and Δ*qim* cells separated and detected by HPLC-FLD (excitation 360 nm, emission 425 nm).

The same glycan compositional analysis performed on the GlcN-phosphate-linked GAC was utilized for the analysis of fully acid-hydrolyzed lipoteichoic acid isolated and purified from QS-ON GAS and the QS-ON-Δ*qim* mutant ^65^. This analysis showed no differences between WT and mutant strains but did reveal an unexpected and unidentified glycan associated with the LTA backbone or linkage unit (Fig. 4B, unidentified peak ∼12 min). These results indicate that the Rgg2/Rgg3 QS system is not modifying the predominant form of GAC or the major glycan profile of LTA.

### The Rgg2/Rgg3 QS system produces cell wall-associated GlcNAc-ribitol

Rush, *et al* determined that roughly 20% of GAC is not deacetylated by PplD and remains coupled to peptidoglycan through GlcNAc-phosphate linkages that are susceptible to mild acids ^57, 66, 67^. Therefore, extraction of sacculi with a mild acid releases GlcNAc-linked GAC and, in theory, releases any other phosphate-linked carbohydrate, such as wall teichoic acids, though GAS is not reported to produce WTA. Sacculi produced from QS-ON cultures of wildtype and the Δ*qim* isogenic mutant were extracted with 25 mM glycine-hydrochloric acid, further purified by anion exchange chromatography, and analyzed by ultra high-performance liquid chromatography-coupled electrospray ionization tandem-MS/MS mass spectrometry (UHPLC-MS/MS), as has been described for the detection of wall teichoic acid monomers (Figs. 4C, S4)^67^. A HexNAc-ribitol (HexNAc-Rbo) species with a *m/z* of 354.14 was identified in the wildtype sample but was absent in carbohydrate extracted from the mutant lacking the *qim* operon (Fig. 5C,D). By comparing this m/z 354 signal to our database of known Gram-positive WTA repeating-unit structures, we found that its retention time aligns with C2-linked GlcNAc-Rbo, and shows a 0.2-min shift relative to the C4-linked GlcNAc-Rbo observed in wall teichoic acids extracted from L. monocytogenes EGDe and 1042, respectively (Fig. S8) ^67^. Glycan compositional analysis, as previously described, for GAC (GlcN-phosphate linked) and LTA was also performed, indicating the extracted carbohydrate sample contained primarily GlcNAc and rhamnose, as was expected for GAC (Fig. 4E) ^63, 64^. No HexNAc monosaccharides other than GlcNAc were detected by compositional analysis; therefore, we assign the HexNAc-Rbo monomeric unit identified by UHPLC-MS analysis as GlcNAc-Rbo, with GlcNAc likely linked at the C2 position of ribitol, as alternative connectivities would result in distinct retention times, as previously demonstrated by Shen *et al* ^67^. These results demonstrate the presence of an uncharacterized GlcNAc-ribitol unit to the cell wall of GAS whose production is dependent on the Rgg2/Rgg3 QS regulated *qim* operon.

**Figure 5.**
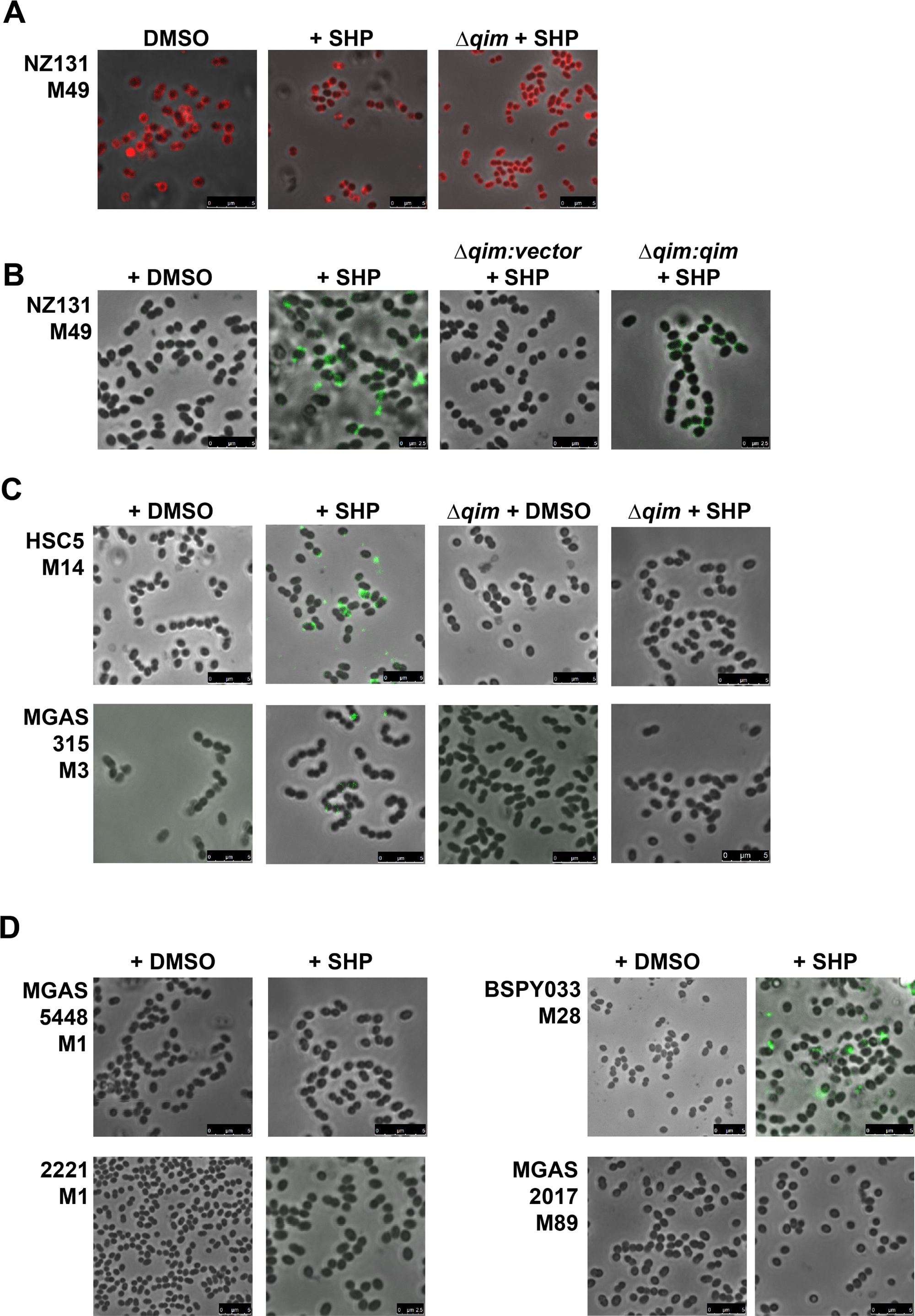
Fluorescent phage proteins detect surface polysaccharides. **(A)** NZ131 (serotype M49) and isogenic Δ*qim* mutant stained with PlyCB-AlexaFluor555 to detect polyrhamnose. 100 nM SHP (QS-ON) or DMSO vehicle (QS-OFF) was added to cultures as indicated. **(A-D)** Bacterial labeling with RBP-13-GFP, which recognizes ribitol phosphate. **(B)** NZ131 (WT), Δ*qim*, and Δ*qim*:*qim* complemented strains cultured without (DMSO) or with 100 nM SHP. **(C)** Serotypes M14 (HSC5) and M3 (MGAS315), and the isogenic qim mutants grown without (DMSO) or with 100 nM SHP. **(D)** Serotypes M1 (5448, 2221), M28 (BSPY033), and M89 (MGAS2017) were grown without (DMSO) or with 100 nM SHP.

To evaluate GAS surface polysaccharides and confirm the presence of the RboP-GlcNAc moiety, we utilized recombinant fluorescent phage proteins that recognize specific glycans. PlyCB recognizes GAS poly-rhamnose^68^ whereas the Φ13 phage receptor binding protein (RBP) binds ribitol-phosphate of staphylococcal wall teichoic acids ^69, 70^. AlexaFluor 555-conjugated PlyCB or GFP-fused RBP-13 were added to GAS cells cultured in the presence or absence of SHP and analyzed by fluorescence microscopy. No differences were observed for recognition of the GAC rhamnose backbone among QS-OFF, -ON and the Δ*qim* mutant (Fig. 5A), indicating that the GAC rhamnose backbone remains unaffected. In contrast, RBP-13-GFP recognition of RboP was seen only when the wildtype strain was induced with SHP (Fig. 5B). The isogenic NZ131 Δ*qim* mutant, even when stimulated with SHP, showed no labeling by RBP-13-GFP, but genetic complementation restored fluorescence. Deletion mutants of the *qim* operon were generated in two other serotypes, M3 (MGAS315) and M14 (HSC5), and both lost SHP-induced RBP-13-GFP recognition (Fig. 5C). Among four other strains tested, we observed RBP-13-GFP binding only for the M28 strain BSPY033 after SHP stimulation, but not for two M1 and one M89 serotypes (Fig. 5D). Taken together, these results validate the presence of RboP on GAS surfaces of multiple serotypes when the Rgg2/3 QS system is induced and display requires the *qim* operon.

## DISCUSSION

It was previously established that the Rgg2/Rgg3 QS-regulated operon *spy49_0450-0460*, renamed here to the **q**uorum-regulated **i**mmunomodulatory **m**odification (*qim*) operon, plays a central role in altering the innate immune response during *in vitro* macrophage infection, and initial characterization of this phenotype highlighted the contact-dependent suppression of the NF-κB response^16, 17^. Here, we confirmed that removal of the operon led to loss of the immunomodulatory phenotype and show that individual genes *qimA*, *qimB*, *qimE*, *qimF*, *qimG*, and *qimH* are required. By placing the operon under a constitutive promoter for complementation, we decoupled its expression from the QS system and demonstrate that these genes alone are sufficient for suppression of the macrophage response independent of other QS-induced changes. Furthermore, our analyses of surface-associated carbohydrates revealed a GlcNAc-ribitol wall teichoic acid (WTA)-like modification not previously described in GAS produced by the *qim* operon.

*In silico* modeling and analysis of the genes within the *qim* operon suggested the operon is involved in the biosynthesis or modification of cell wall-associated carbohydrate or teichoic acids. The only gene in the operon previously shown not to be required for the immunomodulatory phenotype, *qimJ* (*spy49_0460*), was identified as a gene with lipoteichoic acid flippase (LtaA) homology (see supplement Fig. S6) ^17^. Given that LTA is produced by GAS irrespective of QS status, we suspect that *qimJ* is functionally redundant with a housekeeping flippase, possibly *spy49_0419*, which is homologous to LtaA from *Staphylococcus aureus*. It will be interesting to see if the immunosuppressive phenotype is lost when both *qimJ* and *spy49_0419* are disrupted, an experiment that will likely require conditional inhibition due to gene essentiality using technology like CRISPRi ^71^. Nevertheless, analysis of LTA found no obvious differences between QS activity states (ON vs OFF) or in mutants of the *qim* operon, thus we limited our analysis to the LTA glycan composition. However, future studies could look further to elucidate whether alternative linkage patterns are present or whether modifications not detectable by the anthranilic acid (2-AA) method used in our approach are present.

The *qimI* gene was previously characterized as a UDP-glucose dehydrogenase that produces glucuronic acid and shown to be a HasB (*spy49_1806*) paralog able to facilitate moderate levels of hyaluronic acid capsule biosynthesis in a *hasB* mutant strain^20^ (20). We previously established that hyaluronic acid capsule did not impact macrophage suppression and is why HA capsule was not analyzed in the current study^17^. Interestingly, Rahbari *et al* demonstrated an intermediate immunomodulatory phenotype in the macrophage NF-κB reporter assay in the absence of the *qimI* gene, and we infer that, similar to the *qimJ* gene, functional substitution by the redundant gene could be responsible for this.

In addition to having homology with enzymes involved in hyaluronic acid capsule and lipoteichoic acid biosynthesis, operon genes *qimD* and *qimG* are homologous to enzymes involved in the biosynthesis of lipopolysaccharide (LPS) and wall teichoic acids (WTA) in other bacterial species (Fig. S6)^72, 73^. It is less clear how other genes in the operon, namely *qimA* (shikimate dehydrogenase), *qimC* (F420-0:gamma-glutamyl ligase), and *qimE* (SAM synthase), contribute to the biosynthesis of a carbohydrate or cell wall modification. We suspect it to be possible that multiple modifications are produced upon expression of the *qim* operon or that alterations to metabolic processes by the additional enzymes in the operon play a role in the biosynthesis of the immunomodulatory factor. Future work will attempt to characterize these enzymes individually, confirming their putative functions and elucidate the biosynthetic pathway of the modifications described in this study.

Among the most significant outcomes of our analysis was the finding that mild acid extractions of sacculi released a wall teichoic acid-like structure, GlcNAc-ribitol (HexNAc-Rbo), and its presence depended on expression of the *qim* operon that is controlled by the Rgg2/Rgg3 QS system. To the best of our knowledge, GAS has never been reported to produce ribitol-phosphate, a common monomeric unit of WTA in other bacterial species. Although some overlap exists between biosynthesis of WTA and LTA, the LTA of GAS contains only a polyglycerol phosphate backbone. Our attempts at further structure elucidation and linkage analysis of the detected GlcNAc-Rbo moiety by NMR proved challenging as we believe it to be a minor component of the overall GAC structure (or other uncharacterized wall glycan), not easily observed through standard NMR experiments. Interpretation of low intensity NMR signals from polysaccharides often requires specialized experiments and optimization that have thus far been unattainable, but future efforts will attempt to enrich the ribitol-containing carbohydrate to compensate for sensitivity issues with NMR analysis.

The influence of GAS on cultured macrophages clearly indicates that production of cytokines and inflammatory markers are blocked under QS-ON conditions ^16, 17^, but how these activities impact bacterial survival in the context of a functioning immune system is not fully understood. Previously, we demonstrated that mutants of the Rgg2/Rgg3 QS system had a striking impact on colonization of the upper respiratory tract, where a QS-OFF mutant (Δ*rgg2*) was less effective than wildtype or the QS-ON mutant (Δ*rgg3*) in its ability to attach and initiate colonization^74^. To evaluate the contribution that each of the two operons regulated directly by the Rgg2/Rgg3 QS system could have during *in vivo* infection, we utilized a murine skin colonization model in which Wilkening *et al* had found the Rgg2/Rgg3 QS regulated loci to be highly upregulated during GAS infection^58^. Substantial differences in disease severity, indicated by weight change, CFU count, and visual indication of pyoderma and edema, were seen between wildtype and the isogenic *qim* operon mutant; differences between wildtype and the *stcA* mutant were observed to a lesser degree. Together, these results not only indicated that the Rgg2/Rgg3 system to be important in this dermal infection model but also demonstrate the enhanced contribution provided by the *qim* operon in bacterial virulence. How and if disease severity is attributable to macrophage suppression will require a deeper analysis of immune cell signaling, recruitment, and efficacy *in vivo*. As pyodermic exudate was an obvious outcome of wildtype GAS infection, a long-lasting block on inflammatory responses must not be occurring over the entire course of infection. Perhaps instead, down-regulation or delay of the immune response at early stages of microbial exposure provides enough opportunity for GAS to colonize and expand before a vigorous inflammatory response ensues.

The bacterial cell surface is a highly complex and dynamic structure at the interface of host-pathogen interactions, and modifications or alterations to this structure play a critical role in a bacterium’s ability to adapt to and possibly subvert host defenses. Streptococcal glycans and teichoic acids have been shown to inhibit immune signaling pathways. High molecular weight hyaluronic acid of GAS capsule and sialylated capsular polysaccharide of Group B Streptococcus each bind to human Siglec-9 and down-regulates neutrophil activation^75, 76^. And it was shown that an LTA lipid anchor, diglucosyldiacylglycerol (DGDG) inhibits stimulation of Mincle, a c-type lectin receptor expressed on myeloid cells ^36^. We speculate that display of the RboP-GlcNAc moiety on the GAS surface may stimulate an immunoregulatory receptor, such as a siglec or c-type lectin that recruit inhibitory adaptors used by the innate immune system for homeostatic regulation. Attempts to identify this receptor, which we expect to be conserved in mouse and human cells because macrophages of both species display suppression by QS-ON GAS, are ongoing.

Bacterial surface structures not only influence how the host perceives and responds to the pathogen, but they also present a target for host-defense mechanisms as well as attachment sites for bacteriophage^77, 78^. Investigating the advantages as to why GAS has evolved to employ an intercellular signaling (QS) system to regulate what appears to be a significant pathogenic advantage begs the question as to what fitness tradeoffs are gambled if expression of *qim* operon genes were to be constitutively expressed? It is well established that the cell wall binding domain of bacteriophage endolysins rely on surface-associated carbohydrates or wall teichoic acids for attachment, and the production of a GlcNAc-ribitol modification to the surface of GAS cells by the *qim* operon could confer increased bacteriophage susceptibility, and seems likely given the ability of the Φ13-RBP to recognize SHP-induced GAS (Figure 5)^79, 80^. Likewise, changing the electrostatic properties of the cell surface presents both advantages and disadvantages in the context of host defenses. As was seen in the attachment of glycerol-phosphate (GroP) to the GlcNAc side chains of GAC, the modification conferred increased resistance to Zn^2+^ toxicity used by the host immune system to prevent bacterial intracellular survival after phagocytosis by immune cells^43^. However, due to the increased negative charge imparted by the phosphate linkage of GroP to GAC, the bacteria became more susceptible to the human Group IIA phospholipase (hGIIA), a highly cationic protein that binds the bacterial cell surface^43, 81^. Thus, regulating surface structures to match threats of changing situations illustrates the sophistication of bacterial pathogens. The advantage provided by *social* regulation of surface structures (QS regulation) may be driven by a need for conformity among a population, to unify their appearance or coordinate their assault against the host. In this scenario, the QS system does not necessitate a high population density but instead relies on intercellular signaling—even among a small number of bacteria—to coordinate gene expression. The fact that production of SHP pheromones is subject to positive feedback allows for rapid messaging between bacteria to present themselves in a consistent manner that may benefit the group overall. Understanding the benefits afforded to the pathogen by these regulatory systems now offers an opportunity to disrupt bacterial coordinated behaviors as a means of therapeutic intervention.

## MATERIALS AND METHODS

### Bacterial strains and culture conditions

*S. pyogenes* (Group A Streptococcus; GAS) strain NZ131 was used for this study and as the parent strain for all GAS genetic mutants. GAS strains were grown without shaking in Todd-Hewitt medium with 2% (wt/vol) yeast extract (THY, BD Bacto^TM^) broth or on agar plates, or in a chemically-defined medium (CDM) supplemented with 1% (wt/vol) glucose ^82^ Growth conditions were maintained at 37° C with 5% CO_2_. When appropriate antibiotic resistance-based strain selection was accomplished by supplementing the media with erythromycin (Erm), 0.5 µg/mL. For cloning, laboratory *E. coli* strain XL-10 Gold (Agilent) was grown in NZY^+^ broth or Luria-Bertani (LB, BD Difco) according to manufacturer specifications.

General growth conditions for GAS strains were as follows. Strains were streaked onto THY agar and incubated overnight at 37° C + 5% CO_2_. Single colonies were inoculated into sterile THY broth and incubated overnight at 37° C before being pelleted and resuspended in CDM to inoculate at an OD_600_ of ∼ 0.05. Cultures were incubated at 37 °C until reaching an OD_600_ of ∼ 1.0 at which point glycerol was added to a final concentration of 20% and stored at - 80° C in single use aliquots as starter cultures. Starter cultures were routinely used to inoculate CDM at a starting OD_600_ of ∼ 0.05. For QS-ON conditions, cultures were incubated at 37° C for approximately 30 minutes before the addition of SHP-C8 peptide. SHP3-C8 peptide was purchased from Abclonal (Woburn, MA), sequence *DIIIIVGG*, and added to the culture for a final concentration of 100 nM (from 100 µM stock, in DMSO). An equivalent volume of DMSO was used in QS-OFF culture conditions. Cultures were incubated to mid-exponential phase and harvested by centrifugation, washed with sterile PBS, and stored as pellets at -80° C for analysis.

### Mutant strains and plasmid construction

All strains and plasmids used in this study can be found in **Table S1** of the supplemental material, with primers used for plasmid construction in **Table S2**. Cloning was performed using laboratory *E. coli* strain XL-10 Gold (Agilent) with antibiotic selection using erythromycin supplemented LB media at a concentration of 500 µg/mL. For complementation of *qim*, the complementation vector, pJC479, was amplified by PCR in two fragments, then placed into an integrating shuttle vector, p7INT, under the constitutive *syncat* promoter, assembled using the NEBuilder HiFi Assembly kit (New England Biolabs). The vector was then transformed into XL-10 Gold chemically competent *E. coli* per manufacturer instructions with recovery in SOC broth and selection on LB agar with erythromycin. The plasmid was confirmed by whole plasmid sequencing (Plasmidsaurus, Eugene, OR). The complementation vector was then electroporated into GAS strain JCC303 (Δ*qim*) and plated on THY agar supplemented with 0.5 µg/mL of erythromycin for antibiotic selection. Single gene deletions were accomplished by inverse PCR of the complementation vector, omitting one gene, with assembly by using the NEBuilder HiFi Assembly kit or with ligation using T4 DNA ligase (New England Biolabs) after digestion with MluI/DpnI restriction enzymes; the MluI restriction site was added to the 5’ end of inverse PCR primers to facilitate re-ligation. Plasmids were electroporated into GAS strain JCC303 as previously described, and genotypes were confirmed by PCR amplification of genomic DNA isolated from the mutant strains.

### Bioinformatics analysis of the *qim* operon

The *qim* operon genes were analyzed for protein sequence homology using the HHpred server for protein homology detection. The amino acid sequences were uploaded and referenced against the PDB_mmCIF70_12_Aug structural/domain database using all default parameters. This produced a results table of hits with homologous protein structures in the RCSB Protein Data Bank (RCSB PDB). AlphaFold 2.0 was then used to generate predicted monomeric protein structures for each gene in the operon, performed on a HP Z6 workstation with a Xeon Gold 6354 CPU, 192 GB RAM, Nvidia RTX 2080TI GPU and M2 SSD disks, operating on Ubuntu Linux 20.04 with completed databases from 11-01-2021 ^59^. Five models were predicted for each monomer and internally scored using intrinsic model confidence values, with each protein and chain scored using a per residue predicted local-distance difference test (pLDDT). The model with the highest pLDDT value for each prediction was then aligned to the top homologous protein structure identified by HHpred using the super command after visualization in PyMol to yield RMSD values for alignments. Homologous protein structures identified by HHpred were obtained from the RCSB PDB ^60^.

### Chemical extraction of QS-ON GAS cells

GAS was grown as previously described with the Rgg2/Rgg3 QS system induced by the addition of exogenous SHP peptide. Cells were grown to mid-exponential phase and then harvested by centrifugation, washed with PBS, and extracted with chemical solvents covering a range of polarities. Extracts were generated using acetonitrile (ACN):dH_2_O (1:1), acidified ACN:dH_2_O (0.1 M formic acid), basified ACN:dH_2_O (0.1 M sodium hydroxide, NaOH), ethyl acetate (EtOAc), methanol (MeOH):dH_2_O (1:1), and dH_2_O as a cell lysate. All chemicals were purchased from Fisher Scientific (Hampton, NH) as HPLC grade solvents. After extraction, the samples were centrifuged to pellet the cells and the solvent supernatant was removed. Extractions were completed three times and pooled before drying under reduced pressure at 30° C using rotary evaporation (Büchi, New Castle, DE). Dried extracts were reconstituted in DMSO for testing in the macrophage NF-κB reporter assay as described below at a final concentration of 1 mg/mL in 200 µL of macrophage cell culture medium.

### Lipoteichoic acid (LTA) extraction and purification

Frozen culture pellets were thawed, and lipoteichoic acid was extracted and purified using previously described methods ^65^ (59). Pellets were resuspended in dH_2_O for mechanical lysis by French press at 29 psi (2 bar). The cell suspension was then centrifuged at 20,000 x *g* for 20 minutes at 4° C, resuspended in 0.1M Na citrate (pH 4.7) and extracted with an equal volume of *n*-butanol (Fisher Scientific), shaking at 37° C for 30 minutes. Samples were centrifuged at 20,000 x *g* for 30 minutes, 4° C, to facilitate phase separation, and the bottom aqueous layer was removed to a new conical tube. This was centrifuged again at 20,000 x *g* for 40 minutes, 4 °C, transferred to a new conical tube, frozen, and lyophilized. The dried crude LTA extract was resuspended in 5 mL of equilibration buffer (0.1M Na citrate, pH 4.7, 15% *n*-propanol) and purified by hydrophilic interaction chromatography with an octyl Sepharose (HiTrap Octyl FF, Cytiva 17135901; 5 x 1 mL) column using an ÄKTA chromatography system (Cytiva). The column was equilibrated with 6 column volumes (CV) of equilibration buffer, after which the sample was loaded by emptying the sample loop with 10 mL of equilibration buffer. The column was then washed with 5 CV of equilibration buffer. Sample was eluted using a gradient from 100% buffer A (50 mM Na citrate, pH 4.7, 15% *n*-propanol) to 100% buffer B (50 mM Na citrate, pH 4.7, 60% *n*-propanol) over 10 CV, followed by a column wash with 100% buffer B for 4 CV. Re-equilibration of the column was performed between samples with 6 CV of equilibration buffer. Fractions were collected in 2 mL increments, and a flow rate of 5 mL/min was used for all steps. Fractions were assayed for phosphate content as described below and those containing LTA were pooled and dialyzed extensively against dH_2_O with 1000 MWKO dialysis tubing before lyophilization.

### Phosphate assay

Evaluation of the total phosphate content was adapted from protocols described by Draing *et al*., 2006 and Rush *et al*, 2022 ^57, 83^. Briefly, in a 2 mL microcentrifuge tube with secure-lock cap, 100 µL of sample was mixed with 200 µL of washing solution (2M H_2_SO_4_, 0.44M HClO_4_) and heated at 110° C for 2 hours. After cooling to room temperature, 1 mL of reducing solution (3 mM ammonium molybdate, 0.25M Na acetate, 1% ascorbic acid) was added to the sample and incubated for 2 hours at 45° C before measuring the absorbance at 700 nm using a plate reader.

### Isolation of peptidoglycan sacculi

Frozen culture pellets were thawed and resuspended in dH_2_O before being mechanically lysed by French press at a pressure of 29 psi (2 bar) to a final volume of 40 mL. The cell suspension was centrifuged at 20,000 x *g* for 20 minutes and the pellet was resuspended in SM buffer with DNase and RNase A to digest overnight, shaking, at 4° C. After overnight digestion, Proteinase K was added and incubated shaking for 2.5 hours at 37° C. SDS (10%) was added for a final concentration of 4% SDS, and the samples were boiled in a water bath for 1 hour. SDS was removed by 5 rounds of centrifugation at 20,000 x *g* for 20 minutes at 25° C, washing with dH_2_O. The pellet was then transferred to 1.5 mL Eppendorf tubes and stored at -20° C or taken directly for analysis.

Alternatively, peptidoglycan was prepared as described by Kühner *et al*, 2014, stopping before the acid extraction for removal of wall teichoic acids ^84^.

### Estimation of total carbohydrate content

Purified GAS sacculi and whole cells were assayed to estimate total carbohydrate using a modified anthrone assay as described by Rush *et al* ^57^. Briefly, 80 µL of purified sacculi (OD_600_ ∼ 0.3) or whole cells were combined with 320 µL of anthrone reagent (0.2% anthrone in conc. H_2_SO_4_) in capped glass tubes. Cultures were grown under conditions as previously described, when indicated, in the QS-ON state for approximately 3 generations. Samples were heated at 100° C for 10 minutes and then cooled to room temperature to measure UV absorbance at 620 nm. Results were normalized by sacculi/culture optical density.

Carbohydrate content was also evaluated using the phenol-sulfuric acid carbohydrate assay. Briefly, 200 µL of sample was mixed with 200 µL of phenol reagent (5% phenol v/v in dH_2_O) to which 1 mL of concentrated H_2_SO_4_ was rapidly added in capped glass culture tubes. The mixture was allowed to rest for 10 minutes before incubation at 30° C for 20 minutes. Then 300 µL was transferred to 96-well plates and the UV absorbance read at 490 nm.

### Group A Carbohydrate extraction by HONO deamination and purification

GAC was extracted as described by Rush *et al* ^43, 57^. Briefly, GAS sacculi were resuspended in dH_2_O followed by addition of 1 M Na acetate (pH 4.5) to a final concentration of 0.2 M based on the final reaction volume. Over 90 minutes and in three equal volumes, 5 M NaNO_2_ was added for a final concentration of 1.5 M NaNO_2_ with gentle agitation between additions. One molar equivalent of ethanolamine was added to quench the reaction, and the samples were centrifuged for 30 minutes at 16k x *g*. Supernatants were transferred to 3k MWCO spin columns (Amicon Ultra-0.5 mL) and centrifuged at max speed for 13 minutes. The cell pellet was resuspended in 200 µL dH_2_O, centrifuged as before, and the supernatant added to the spin column followed by a second round of centrifugation. The cell pellet was washed again with 400 µL of dH_2_O, centrifuged, and again the supernatant added to the spin column and concentrated. The retentate was desalted by four rounds of dilution with 400 µL of dH_2_O and then eluted by vortexing with 50 µL of dH_2_O followed by inversion of the spin column and centrifugation into a clean Eppendorf tube. Sample recovery was repeated with an additional 300 µL of dH_2_O before purification by size exclusion chromatography using a Superdex 200 Increase 10/300 GL column on an ÄKTA chromatography system (Cytiva). Samples were eluted with dH_2_O over 1.5 column volumes (CV) with a flow rate of 0.75 mL/min while collecting fractions in 0.5 mL increments and monitoring UV absorbance at 205 nm and 212 nm. Fractions were evaluated for carbohydrate presence by phenol-sulfuric acid assay as described above and carbohydrate containing fractions were pooled for anion exchange chromatography (AE) using a HiTrap DEAE FF column (5 mL; GE Healthcare, Glattbrugg, Switzerland). Samples were loaded onto the DEAE column and after an isocratic hold of 4 CV with 10 mM Tris-HCl (pH 7.5), a gradient method was used from 0-1 M NaCl in 10 mM Tris-HCl (pH 7.5) over 10 column volumes with a 3 mL/min flow rate. Fractions were collected in 2 mL increments and UV absorbance was monitored at 205 nm and 212 nm. The fractions were pooled as indicated by the UV chromatogram for non-retained and retained species and desalted against dH_2_O in 1k MWCO dialysis tubing over 3 days. Following dialysis, the samples were lyophilized before further analyses were conducted.

### Mild acid (glycine-HCl) carbohydrate extraction and purification

Isolated GAS sacculi were extracted in 25 mM glycine-HCl for 10 minutes at 100° C in a hot water bath as described for the extraction of wall teichoic acids ^67^. The samples were then centrifuged at 30,000 x *g* for 30 minutes, the pellet was re-extracted, centrifuged as before, and supernatants were pooled before extensive dialysis against dH_2_O with 1 kDa MWCO dialysis tubing, followed by lyophilization. Dried samples were resuspended in 5 mL dH_2_O and purified by anion exchange chromatography as performed for the final purification of GAC described above.

### Monomer HF digestion analysis of mild-acid extracts by UPLC-MS/MS

Carbohydrate samples were treated to hydrolyze all phosphodiester bonds for structural and compositional analysis by UPLC-MS/MS ^67, 85^. Samples were digested with 100 µL of 48% hydrofluoric acid (HF) for 16 hours at 4 °C in an ice water bath and then lyophilized under NaOH pellets to neutralize HF vapors before being reconstituted in dH_2_O for analysis. Monomer UPLC-MS/MS analysis was conducted as previously described by Shen *et al.* for the analysis of depolymerized wall teichoic acid monomers ^67^. Wall teichoic acid with a known C4-linked GlcNAc-RboP monomer from the *Listeria monocytogenes* strain 1042 was analyzed by the same methods and used as a standard for *m/z* and retention time matching ^67^.

### 2AA-Labeled Glycan Compositional Analysis

Glycan compositional analysis was performed based on methods described by Anumula, *et al* ^63, 64^. Carbohydrate samples were digested for complete acid hydrolysis with 20% trifluoroacetic acid (TFA) at 98° C for 5 hours or overnight while shaking at 800 rpm before being dried by centrifugal evaporation under reduced pressure at 60° C.

For labelling, the dried acid hydrolyzed samples were resuspended in 1% sodium acetate (NaOAc) for 30 minutes with periodic vortexing before adding anthranilic acid (2-AA) reagent. 2-AA reagent was prepared by mixing equal parts of cyanoborohydride (CBN) solution (40 mg CBN in 1 mL NBM solution (2.4% NaOAc, 2% boric acid in methanol)) and 2-AA solution (60 mg 2-anthranilic acid in 1 mL NBM solution; 220 mM final concentration). The reaction mixture was incubated at 80° C, shaking (750 rpm) for 45 minutes, covered with aluminum foil. Samples were then cooled to ambient temperature and stored at 4° C before analysis.

For HPLC analysis, concentrated samples were diluted with mobile phase A (0.3% 1-amino butane, 0.5% phosphoric acid, 1% tetrahydrofuran (THF) in dH_2_O), mixed well, and centrifuged at 20,000 x *g* for 20 minutes. Monosaccharides were separated over a YMC-Pack ODS-A (5 um particle size, 12 nm pore size, 4.6 x 150 mm) column with a flow rate of 1.5 mL/min with fluorescence detection (Ex. 360 nm, Em. 425 nm). Sample was eluted using an isocratic method of 94% mobile phase A and 6% mobile phase B (50% mobile phase A, 50% acetonitrile) for 20 minutes, followed by a column wash with 100% mobile phase B for 1 minute before re-equilibrating with the starting conditions. Monosaccharide standards were analyzed for comparison, prepared as described above by complete acid hydrolysis and labelling of 100 nanomoles of each monosaccharide.

### NMR analysis

The purified GAC polymers of wildtype GAS strain NZ131 and Δ*qim* strains were analyzed by NMR spectroscopy for structural elucidation. A series of experiments, including ^1^H proton and ^1^H-^13^C-HSQC were acquired in D_2_O at 25 °C. NMR spectra were recorded on a Bruker Avance III HD 600 MHz spectrometer equipped with a Prodigy triple-resonance probe and a Bruker Avance III HD 500MHz spectrometer equipped with a BBFO broadband probe. Spectra were analyzed using MestRe Nova 14.

### NF-κB reporter assay

Macrophages expressing a secreted embryonic alkaline phosphatase (SEAP) reporter construct inducible by NF-κB (RAW-Blue^TM^, InvivoGen) were used to evaluate the immunomodulatory potential of GAS strains and purified cell components or chemical extracts as described by Rahbari *et al* with described modifications ^17^. Macrophages were grown in Dulbecco modified Eagle medium (DMEM) (Gibco) with 10% fetal bovine serum (FBS) (Gemini, BenchMark) and maintained at 37° C with 5% CO_2_. Cells were seeded into tissue culture treated T25 flasks (Greiner Bio-One) and grown to approx. 70% confluence before being passaged and plated for testing. The cells were washed with PBS, incubated at 37° C with 0.05% trypsin-0.53 mM EDTA (Corning) for 5 minutes, and then an equal volume of DMEM was added to inactivate the trypsin. Macrophages were counted and seeded into a tissue culture treated 96-well plate (Corning) at 2.5 × 10^4^ cells/well to be used the following day for testing. For bacterial assays, fresh DMEM was added to the macrophage reporter cells with GAS strains added at a multiplicity of infection (MOI) of 40:1. To equilibrate the infection, the plate was centrifuged for 5 minutes followed by a 30-minute incubation at 37° C with 5% CO_2_. At this point the media was removed and replaced with DMEM supplemented with 100 µg/mL gentamicin and 100 ng/mL lipopolysaccharide (LPS, Sigma). At either 8 hours post infection (hpi) or 18 hpi, the cell culture supernatants were collected and combined with the QUANTI-Blue reagent (InvivoGen) and incubated at 37° C in a flat bottom 96-well plate before measuring the absorbance at 625 nm (Synergy HTX microplate reader, BioTek). Results were normalized to LPS when appropriate, and QS-ON values were normalized to QS-OFF conditions when evaluating the QS-dependent immunomodulatory potential of a given GAS strain.

Chemical extracts and purified cell components were tested in the macrophage NF-κB reporter assay following a similar protocol; instead of infecting with bacteria, the extract or cellular component was supplemented to the macrophage cell culture medium along with gentamicin and LPS as previously indicated. QS-ON GAS extracts were reconstituted in DMSO and tested at 1 mg/mL in DMEM supplemented with gentamicin and LPS. Concentrations of DMSO in DMEM did not exceed 1% total culture volume.

### Carbohydrate binding by Fluorescent Phage Proteins

Cultures of GAS grown to mid-log phase in the absence or presence of 100 nM SHP were washed in PBS and ∼1×10^5^ CFU in 50 ul were incubated with 5 μg/ml of either PlyCB-AlexaFluor555 ^68^ or RBP-13-GFP ^69^ for 30 minutes. Cells were washed with PBS prior to mounting on glass slides and imaged by confocal laser scanning microscopy.

### Mouse Colonization

Mice were housed in the BSL2 facilities provided by the University of Colorado School of Medicine. All experiments were conducted under approved IACUC protocols and as described by Wilkening *et al* ^58^. Food and water were provided *ad libitum* during experiments, with supplementary dry feed provided on the floor of cages during extended infection experiments. Male C57 BL6 mice obtained from Jackson Labs were used for all experiments, timed such that all mice were 8 weeks old at start of experiments. Mice were allowed to acclimate for at least 1 week in the vivarium prior to experimentation. 1 day prior to inoculation, the dorsum of the mice was shaved using battery operated trimmers. Residual hair was removed with Nair and mice were allowed to recover overnight before inoculation.

Prior to experimentation, bacterial starter cultures were prepared by growing all strains to mid-log phase. Sterile glycerol was added to a final concentration of 20% and 2 ml aliquots were frozen at -80° C. On the day of inoculation, starter cultures were used to produce input bacterial cultures. These cultures were grown to mid-log phase and diluted to attain approximately 10^8^ CFU/200 µL inoculum. 200 µL of the inoculum were applied to sterile band-aid pads and affixed to the backs of depilated mice under isoflurane anesthesia. Band-aids were then covered with a Tegaderm occlusive dressing (3M) and secured with a second band-aid. At each time point, the dressing was removed, mice were weighed and photographed. In addition, on days 1, 5, and 7 mice were swabbed with a pre-wetted cotton swab. Swabs were placed in sterile PBS, vortexed for 30 seconds each, serially diluted and plated on selective GAS CHROMagar^TM^ (Paris, France) medium to assess bacterial burden after approximately 24 hours of incubation at 37° C + 5% CO_2_. Following all interventions, a new dressing was applied and mice were allowed to recover from anesthesia. Mice were observed throughout all experiments for signs of discomfort or illness. Any mouse with greater than 25% weight loss from pre-experimental weight were declared moribund and sacrificed. At 7 days post-infection, mice were sacrificed using CO_2_ euthanasia.

## Author Contributions

Caleb M. Anderson – experimental design, manuscript, performed experiments and data analysis

Reid Wilkening and Alex Horswill – experimental design and performed animal studies

Samy Boulos – performed mass spectrometry experiments (UHPLC-MS/MS, monomer analysis)

Timothy Keys – experimental design (glycan compositional analysis)

Marc-Olivier Ebert – NMR experimental design, data acquisition

Léa V. Zinsli – AlphaFold structure predictions

Janes Krusche –constructed fluorescent protein expression plasmid

Sam Feldstein – performed cell culturing and inflammatory response experiments

Jennifer C. Chang – experimental design, constructed plasmids and mutants

Andreas Peschel – experimental design of phage receptor binding protein

Martin J. Loessner – funding acquisition, conceptualization

Yang Shen – experimental design, data analysis

Michael J. Federle – manuscript, funding acquisition, experimental design, and data analysis

## Data Availability

Supplementary data and high-resolution images of mouse infections are available here: https: https://figshare.com/s/7b0cb627ed84429172fe

## Acknowledgements

We are grateful to Kate Rahbari, Natalia Korotkova, Dara Kiani and members of the Federle and Loessner labs for suggestions and discussions. Efforts were supported by NIH NIAID grants AI091779 and AI162679 to MJF, and AI153185 and AI162964 to Dr. Horswill. CMA was supported by the DOD SMART Scholarship-for-Service. RVW received salary support from NIH NIAMS grant T32AR007534-36 and the Section of Pediatric Critical Care Medicine in the Department of Pediatrics at the University of Colorado School of Medicine. Mouse graphic in Figure 2 is Adobe stock image 135214677.

## REFERENCES

1. In: Ferretti JJ, Stevens DL, Fischetti VA, editors. Streptococcus pyogenes: Basic Biology to Clinical Manifestations. 2nd ed. Oklahoma City (OK) 2022.

2. Castro SA, Dorfmueller HC. A brief review on Group A Streptococcus pathogenesis and vaccine development. R Soc Open Sci. 2021;8(3):201991.

3. Ralph AP, Carapetis JR. Group a streptococcal diseases and their global burden. Host-pathogen interactions in streptococcal diseases. 2012:1–27.

4. Sims Sanyahumbi A, Colquhoun S, Wyber R, Carapetis JR. Global Disease Burden of Group A Streptococcus. In: Ferretti JJ, Stevens DL, Fischetti VA, editors. Streptococcus pyogenes: Basic Biology to Clinical Manifestations. Oklahoma City (OK)2016.

5. Carapetis JR, Steer AC, Mulholland EK, Weber M. The global burden of group A streptococcal diseases. Lancet Infect Dis. 2005;5(11):685–94.

6. Rogers S, Commons R, Danchin MH, Selvaraj G, Kelpie L, Curtis N, et al. Strain prevalence, rather than innate virulence potential, is the major factor responsible for an increase in serious group A streptococcus infections. The Journal of infectious diseases. 2007;195(11):1625–33.

7. Organization WH. Increased incidence of scarlet fever and invasive group A Streptococcus infection—multi-country. World Health Organization. 2022.

8. Walker MJ, Barnett TC, McArthur JD, Cole JN, Gillen CM, Henningham A, et al. Disease manifestations and pathogenic mechanisms of Group A Streptococcus. Clin Microbiol Rev. 2014;27(2):264–301.

9. Miller MB, Bassler BL. Quorum sensing in bacteria. Annu Rev Microbiol. 2001;55:165–99.

10. Jimenez JC, Federle MJ. Quorum sensing in group A Streptococcus. Front Cell Infect Microbiol. 2014;4:127.

11. Chang JC, LaSarre B, Jimenez JC, Aggarwal C, Federle MJ. Two group A streptococcal peptide pheromones act through opposing Rgg regulators to control biofilm development. PLoS Pathog. 2011;7(8):e1002190.

12. Aggarwal C, Jimenez JC, Nanavati D, Federle MJ. Multiple length peptide-pheromone variants produced by Streptococcus pyogenes directly bind Rgg proteins to confer transcriptional regulation. J Biol Chem. 2014;289(32):22427–36.

13. Sulavik MC, Tardif G, Clewell DB. Identification of a gene, rgg, which regulates expression of glucosyltransferase and influences the Spp phenotype of Streptococcus gordonii Challis. J Bacteriol. 1992;174(11):3577–86.

14. Lasarre B, Aggarwal C, Federle MJ. Antagonistic Rgg regulators mediate quorum sensing via competitive DNA binding in Streptococcus pyogenes. MBio. 2013;3(6).

15. Lasarre B, Chang JC, Federle MJ. Redundant group a streptococcus signaling peptides exhibit unique activation potentials. J Bacteriol. 2013;195(18):4310–8.

16. Feldstein SF, Rahbari KM, Leonardo TR, Alvernaz SA, Federle MJ. Suppressed macrophage response to quorum-sensing-active Streptococcus pyogenes occurs at the level of the nucleus. bioRxiv. 2025.

17. Rahbari KM, Chang JC, Federle MJ. A Streptococcus Quorum Sensing System Enables Suppression of Innate Immunity. mBio. 2021;12(3).

18. Gogos A, Jimenez JC, Chang JC, Wilkening RV, Federle MJ. A Quorum Sensing-Regulated Protein Binds Cell Wall Components and Enhances Lysozyme Resistance in Streptococcus pyogenes. J Bacteriol. 2018;200(11).

19. Chen IA, Chu K, Palaniappan K, Ratner A, Huang J, Huntemann M, et al. The IMG/M data management and analysis system v.7: content updates and new features. Nucleic Acids Res. 2023;51(D1):D723–D32.

20. Cole JN, Aziz RK, Kuipers K, Timmer AM, Nizet V, van Sorge NM. A conserved UDP-glucose dehydrogenase encoded outside the hasABC operon contributes to capsule biogenesis in group A Streptococcus. J Bacteriol. 2012;194(22):6154–61.

21. Kizy AE, Neely MN. First Streptococcus pyogenes signature-tagged mutagenesis screen identifies novel virulence determinants. Infect Immun. 2009;77(5):1854–65.

22. Chang JC, Wilkening RV, Rahbari KM, Federle MJ. Quorum sensing regulation of a major facilitator superfamily transporter affects multiple streptococcal virulence factors. Journal of Bacteriology. 2022;204(9):e00176–22.

23. Henningham A, Yamaguchi M, Aziz RK, Kuipers K, Buffalo CZ, Dahesh S, et al. Mutual exclusivity of hyaluronan and hyaluronidase in invasive group A Streptococcus. J Biol Chem. 2014;289(46):32303–15.

24. Hurst JR, Shannon BA, Craig HC, Rishi A, Tuffs SW, McCormick JK. The Streptococcus pyogenes hyaluronic acid capsule promotes experimental nasal and skin infection by preventing neutrophil-mediated clearance. PLoS Pathog. 2022;18(11):e1011013.

25. Krause RM, Mc CM. Variation in the group-specific carbohydrate of group C hemolytic Streptococci. J Exp Med. 1962;116(2):131–40.

26. Lancefield RC. A Serological Differentiation of Human and Other Groups of Hemolytic Streptococci. J Exp Med. 1933;57(4):571–95.

27. Leon O, Panos C. An electron microscope study of kidney basement membrane changes in the mouse by lipoteichoic acid from Streptococcus pyogenes. Can J Microbiol. 1987;33(8):709–17.

28. Mc CM. Further studies on the chemical basis for serological specificity of Group A streptococcal carbohydrate. J Exp Med. 1958;108(3):311–23.

29. Mc CM, Lancefield RC. Variation in the group-specific carbohydrate of group A streptococci. I. Immunochemical studies on the carbohydrates of variant strains. J Exp Med. 1955;102(1):11–28.

30. McCarty M. Variation in the group-specific carbohydrate of group A streptococci. II. Studies on the chemical basis for serological specificity of the carbohydrates. J Exp Med. 1956;104(5):629–43.

31. Slabyj BM, Panos C. Teichoic acid of a stabilized L-form of Streptococcus pyogenes. J Bacteriol. 1973;114(3):934–42.

32. Slabyj BM, Panos C. Membrane lipoteichoic acid of Streptococcus pyogenes and its stabilized L-form and the effect of two antibiotics upon its cellular content. J Bacteriol. 1976;127(2):855–62.

33. Kendall FE, Heidelberger M, Dawson MH. A Serologieally Inactive Polysaccharide Elaborated by Mucoid Strains of Group A Hemolytie Streptococcus. 1937.

34. Stollerman GH, Dale JB. The importance of the group a streptococcus capsule in the pathogenesis of human infections: a historical perspective. Clin Infect Dis. 2008;46(7):1038–45.

35. van Sorge NM, Cole JN, Kuipers K, Henningham A, Aziz RK, Kasirer-Friede A, et al. The classical lancefield antigen of group a Streptococcus is a virulence determinant with implications for vaccine design. Cell Host Microbe. 2014;15(6):729–40.

36. Imai T, Matsumura T, Mayer-Lambertz S, Wells CA, Ishikawa E, Butcher SK, et al. Lipoteichoic acid anchor triggers Mincle to drive protective immunity against invasive group A Streptococcus infection. Proc Natl Acad Sci U S A. 2018;115(45):E10662–E71.

37. Neuhaus FC, Baddiley J. A continuum of anionic charge: structures and functions of D-alanyl-teichoic acids in gram-positive bacteria. Microbiol Mol Biol Rev. 2003;67(4):686–723.

38. Neuhaus FC, Heaton MP, Debabov DV, Zhang Q. The dlt operon in the biosynthesis of D-alanyl-lipoteichoic acid in Lactobacillus casei. Microb Drug Resist. 1996;2(1):77–84.

39. Mistou MY, Sutcliffe IC, van Sorge NM. Bacterial glycobiology: rhamnose-containing cell wall polysaccharides in Gram-positive bacteria. FEMS Microbiol Rev. 2016;40(4):464–79.

40. Sutcliffe IC, Black GW, Harrington DJ. Bioinformatic insights into the biosynthesis of the Group B carbohydrate in Streptococcus agalactiae. Microbiology (Reading). 2008;154(Pt 5):1354–63.

41. Coligan JE, Kindt TJ, Krause RM. Structure of the streptococcal groups A, A-variant and C carbohydrates. Immunochemistry. 1978;15(10-11):755–60.

42. Huang DH, Rama Krishna N, Pritchard DG. Characterization of the group A streptococcal polysaccharide by two-dimensional 1H-nuclear-magnetic-resonance spectroscopy. Carbohydr Res. 1986;155:193–9.

43. Edgar RJ, van Hensbergen VP, Ruda A, Turner AG, Deng P, Le Breton Y, et al. Discovery of glycerol phosphate modification on streptococcal rhamnose polysaccharides. Nat Chem Biol. 2019;15(5):463–71.

44. Atilano ML, Yates J, Glittenberg M, Filipe SR, Ligoxygakis P. Wall teichoic acids of Staphylococcus aureus limit recognition by the drosophila peptidoglycan recognition protein-SA to promote pathogenicity. PLoS Pathog. 2011;7(12):e1002421.

45. Brown S, Santa Maria JP, Jr., Walker S. Wall teichoic acids of gram-positive bacteria. Annu Rev Microbiol. 2013;67:313–36.

46. Diacovich L, Gorvel JP. Bacterial manipulation of innate immunity to promote infection. Nat Rev Microbiol. 2010;8(2):117–28.

47. Hasty DL, Meron-Sudai S, Cox KH, Nagorna T, Ruiz-Bustos E, Losi E, et al. Monocyte and macrophage activation by lipoteichoic Acid is independent of alanine and is potentiated by hemoglobin. J Immunol. 2006;176(9):5567–76.

48. Kojima N, Kojima S, Hosokawa S, Oda Y, Zenke D, Toura Y, et al. Wall teichoic acid-dependent phagocytosis of intact cell walls of Lactiplantibacillus plantarum elicits IL-12 secretion from macrophages. Front Microbiol. 2022;13:986396.

49. Rockel C, Hartung T. Systematic review of membrane components of gram-positive bacteria responsible as pyrogens for inducing human monocyte/macrophage cytokine release. Front Pharmacol. 2012;3:56.

50. Shiratsuchi A, Shimizu K, Watanabe I, Hashimoto Y, Kurokawa K, Razanajatovo IM, et al. Auxiliary role for D-alanylated wall teichoic acid in Toll-like receptor 2-mediated survival of Staphylococcus aureus in macrophages. Immunology. 2010;129(2):268–77.

51. Swoboda JG, Campbell J, Meredith TC, Walker S. Wall teichoic acid function, biosynthesis, and inhibition. Chembiochem. 2010;11(1):35–45.

52. Wilkening RV, Chang JC, Federle MJ. PepO, a CovRS-controlled endopeptidase, disrupts Streptococcus pyogenes quorum sensing. Mol Microbiol. 2016;99(1):71–87.

53. Bera A, Biswas R, Herbert S, Gotz F. The presence of peptidoglycan O-acetyltransferase in various staphylococcal species correlates with lysozyme resistance and pathogenicity. Infect Immun. 2006;74(8):4598–604.

54. Bera A, Herbert S, Jakob A, Vollmer W, Gotz F. Why are pathogenic staphylococci so lysozyme resistant? The peptidoglycan O-acetyltransferase OatA is the major determinant for lysozyme resistance of Staphylococcus aureus. Mol Microbiol. 2005;55(3):778–87.

55. Davis KM, Weiser JN. Modifications to the peptidoglycan backbone help bacteria to establish infection. Infect Immun. 2011;79(2):562–70.

56. Vollmer W, Tomasz A. The pgdA gene encodes for a peptidoglycan N-acetylglucosamine deacetylase in Streptococcus pneumoniae. J Biol Chem. 2000;275(27):20496–501.

57. Rush JS, Parajuli P, Ruda A, Li J, Pohane AA, Zamakhaeva S, et al. PplD is a de-N-acetylase of the cell wall linkage unit of streptococcal rhamnopolysaccharides. Nat Commun. 2022;13(1):590.

58. Wilkening RV, Langouet-Astrie C, Severn MM, Federle MJ, Horswill AR. Identifying genetic determinants of Streptococcus pyogenes-host interactions in a murine intact skin infection model. Cell Rep. 2023;42(11):113332.

59. Jumper J, Evans R, Pritzel A, Green T, Figurnov M, Ronneberger O, et al. Highly accurate protein structure prediction with AlphaFold. Nature. 2021;596(7873):583–9.

60. Zimmermann L, Stephens A, Nam SZ, Rau D, Kubler J, Lozajic M, et al. A Completely Reimplemented MPI Bioinformatics Toolkit with a New HHpred Server at its Core. J Mol Biol. 2018;430(15):2237–43.

61. Gabler F, Nam SZ, Till S, Mirdita M, Steinegger M, Soding J, et al. Protein Sequence Analysis Using the MPI Bioinformatics Toolkit. Curr Protoc Bioinformatics. 2020;72(1):e108.

62. Dreywood R. Qualitative test for carbohydrate material. Industrial & Engineering Chemistry Analytical Edition. 1946;18(8):499-.

63. Anumula KR. Quantitative determination of monosaccharides in glycoproteins by high-performance liquid chromatography with highly sensitive fluorescence detection. Anal Biochem. 1994;220(2):275–83.

64. Anumula KR, Dhume ST. High resolution and high sensitivity methods for oligosaccharide mapping and characterization by normal phase high performance liquid chromatography following derivatization with highly fluorescent anthranilic acid. Glycobiology. 1998;8(7):685–94.

65. Kho K, Meredith TC. Extraction and Analysis of Bacterial Teichoic Acids. Bio Protoc. 2018;8(21):e3078.

66. Eugster MR, Haug MC, Huwiler SG, Loessner MJ. The cell wall binding domain of Listeria bacteriophage endolysin PlyP35 recognizes terminal GlcNAc residues in cell wall teichoic acid. Mol Microbiol. 2011;81(6):1419–32.

67. Shen Y, Boulos S, Sumrall E, Gerber B, Julian-Rodero A, Eugster MR, et al. Structural and functional diversity in Listeria cell wall teichoic acids. J Biol Chem. 2017;292(43):17832–44.

68. Shen Y, Barros M, Vennemann T, Gallagher DT, Yin Y, Linden SB, et al. A bacteriophage endolysin that eliminates intracellular streptococci. Elife. 2016;5.

69. Krusche J, Beck C, Lehmann E, Gerlach D, Daiber E, Mayer C, et al. Characterization and host range prediction of Staphylococcus aureus phages through receptor-binding protein analysis. Cell Rep. 2025;44(3):115369.

70. Krusche J, Beck C, Lehmann E, Gerlach D, Wolz C, Peschel A. Systematic classification of phage receptor-binding proteins predicts surface glycopolymer structure in Staphylococcus pathogens. bioRxiv. 2024:2024.03. 04.583386.

71. Bjanes E, Stream A, Janssen AB, Gibson PS, Bravo AM, Dahesh S, et al. An efficient in vivo-inducible CRISPR interference system for group A Streptococcus genetic analysis and pathogenesis studies. mBio. 2024;15(8):e0084024.

72. Westman EL, McNally DJ, Charchoglyan A, Brewer D, Field RA, Lam JS. Characterization of WbpB, WbpE, and WbpD and reconstitution of a pathway for the biosynthesis of UDP-2,3-diacetamido-2,3-dideoxy-D-mannuronic acid in Pseudomonas aeruginosa. J Biol Chem. 2009;284(18):11854–62.

73. Sobhanifar S, Worrall LJ, King DT, Wasney GA, Baumann L, Gale RT, et al. Structure and Mechanism of Staphylococcus aureus TarS, the Wall Teichoic Acid beta-glycosyltransferase Involved in Methicillin Resistance. PLoS Pathog. 2016;12(12):e1006067.

74. Gogos A, Federle MJ. Colonization of the Murine Oropharynx by Streptococcus pyogenes Is Governed by the Rgg2/3 Quorum Sensing System. Infect Immun. 2020;88(10).

75. Chang YC, Olson J, Beasley FC, Tung C, Zhang J, Crocker PR, et al. Group B Streptococcus engages an inhibitory Siglec through sialic acid mimicry to blunt innate immune and inflammatory responses in vivo. PLoS Pathog. 2014;10(1):e1003846.

76. Secundino I, Lizcano A, Roupe KM, Wang X, Cole JN, Olson J, et al. Host and pathogen hyaluronan signal through human siglec-9 to suppress neutrophil activation. J Mol Med (Berl). 2016;94(2):219–33.

77. King H, Ajay Castro S, Pohane AA, Scholte CM, Fischetti VA, Korotkova N, et al. Molecular basis for recognition of the Group A Carbohydrate backbone by the PlyC streptococcal bacteriophage endolysin. Biochem J. 2021;478(12):2385–97.

78. Nelson D, Schuch R, Chahales P, Zhu S, Fischetti VA. PlyC: a multimeric bacteriophage lysin. Proc Natl Acad Sci U S A. 2006;103(28):10765–70.

79. Sumrall ET, Shen Y, Keller AP, Rismondo J, Pavlou M, Eugster MR, et al. Phage resistance at the cost of virulence: Listeria monocytogenes serovar 4b requires galactosylated teichoic acids for InlB-mediated invasion. PLoS Pathog. 2019;15(10):e1008032.

80. Shen Y, Kalograiaki I, Prunotto A, Dunne M, Boulos S, Taylor NMI, et al. Structural basis for recognition of bacterial cell wall teichoic acid by pseudo-symmetric SH3b-like repeats of a viral peptidoglycan hydrolase. Chem Sci. 2020;12(2):576–89.

81. Djoko KY, Ong CL, Walker MJ, McEwan AG. The Role of Copper and Zinc Toxicity in Innate Immune Defense against Bacterial Pathogens. J Biol Chem. 2015;290(31):18954–61.

82. van de Rijn I, Kessler RE. Growth characteristics of group A streptococci in a new chemically defined medium. Infect Immun. 1980;27(2):444–8.

83. Boveri M, Kinsner A, Berezowski V, Lenfant AM, Draing C, Cecchelli R, et al. Highly purified lipoteichoic acid from gram-positive bacteria induces in vitro blood-brain barrier disruption through glia activation: role of pro-inflammatory cytokines and nitric oxide. Neuroscience. 2006;137(4):1193–209.

84. Kuhner D, Stahl M, Demircioglu DD, Bertsche U. From cells to muropeptide structures in 24 h: peptidoglycan mapping by UPLC-MS. Sci Rep. 2014;4:7494.

85. Ip CC, Manam V, Hepler R, Hennessey JP, Jr. Carbohydrate composition analysis of bacterial polysaccharides: optimized acid hydrolysis conditions for HPAEC-PAD analysis. Anal Biochem. 1992;201(2):343–9.

